# Sustained neural activity correlates with rapid perceptual learning of auditory patterns

**DOI:** 10.1101/2020.09.13.295568

**Authors:** Björn Herrmann, Kurdo Araz, Ingrid S. Johnsrude

**Affiliations:** Rotman Research Institute, Baycrest, M6A 2E1, North York, ON, Canada; Department of Psychology, University of Toronto, M5S 1A1, Toronto, ON, Canada; Department of Psychology, University of Western Ontario, N6A 3K7, London, ON, Canada; School of Communication Sciences & Disorders, University of Western Ontario, N6A 5B7, London, ON, Canada

**Keywords:** Electroencephalography, perceptual learning, memory, regular pattern, sound, auditory perception

## Abstract

Repeating structures forming regular patterns are common in sounds. Learning such patterns may enable accurate perceptual organization. In five experiments, we investigated the behavioral and neural signatures of rapid perceptual learning of regular sound patterns. We show that recurring (compared to novel) patterns are detected more quickly and increase sensitivity to pattern deviations and to the temporal order of pattern onset relative to a visual stimulus. Sustained neural activity reflected perceptual learning in two ways. Firstly, sustained activity increased earlier for recurring than novel patterns when participants attended to sounds, but not when they ignored them; this earlier increase mirrored the rapid perceptual learning we observed behaviorally. Secondly, the magnitude of sustained activity was generally lower for recurring than novel patterns, but only for trials later in the experiment, and independent of whether participants attended to or ignored sounds. The late manifestation of sustained activity reduction suggests that it is not directly related to rapid perceptual learning, but to a mechanism that does not require attention to sound. In sum, we demonstrate that the latency of sustained activity reflects rapid perceptual learning of auditory patterns, while the magnitude may reflect a result of learning, such as better prediction of learned auditory patterns.

## Introduction

Natural sounds such as speech and music are rich in structured amplitude and frequency motifs that recur over time – here referred to as regular patterns (Rosen, 1992; Topbas et al., 2012; Broze and Huron, 2013). Sensitivity to regular patterns is thought to optimize auditory perception (Smith and Lewicki, 2006; Kluender et al., 2013) by enabling, for example, segregation of sound streams (Snyder and Alain, 2007; Bendixen, 2014), detection of acoustic changes (Schröger, 2005; Winkler et al., 2009; Herrmann et al., 2020), and recognition and prediction of sounds (Jones and Boltz, 1989; Henry and Herrmann, 2014; Nobre and van Ede, 2018). Learning of regular patterns may also benefit perception, for example, by increasing detection sensitivity and reducing detection time of recognizable sounds (Agus et al., 2010; Bianco et al., 2020). The current study investigates neural markers related to perceptual learning of regular auditory patterns.

Sustained neural activity is a signature associated with the processing of regular patterns in sound. Sustained activity increases shortly after the onset of a regular pattern and has been observed for a variety of patterns, including repeated sequences of tone pips (Barascud et al., 2016; Southwell et al., 2017; Herrmann and Johnsrude, 2018b; Southwell and Chait, 2018), coherent chord patterns (Teki et al., 2016), and periodic amplitude and frequency modulations (Gutschalk et al., 2002; Ross et al., 2002; Herrmann and Johnsrude, 2018b; Herrmann et al., 2019). This activity appears to be generated in the auditory cortex (Pantev et al., 1994; Pantev et al., 1996; Gutschalk et al., 2002; Barascud et al., 2016), and possibly also the parietal cortex, frontal cortex, and hippocampus (Tiitinen et al., 2012; Barascud et al., 2016; Teki et al., 2016).

It is not clear whether sustained neural activity in response to regular patterns reflects a stable, stimulus-driven representation of an auditory pattern ‘gestalt’ or ‘object’ (Shinn-Cunningham, 2008), or whether pattern-related sustained activity can also be modulated by experience with a given pattern. We can distinguish these possibilities by contrasting sustained activity to regular patterns that sporadically recur over trials with sustained activity to regular patterns that are trial-unique (Luo et al., 2013; Andrillon et al., 2015; Bianco et al., 2020). If sustained activity is driven entirely by the regularity of the pattern, we would expect it to be identical regardless of whether a regular pattern is trial-unique or recurring. If, however, sustained activity is different for recurring patterns compared to unique patterns, then it may also reflect processes related to perceptual learning and is more cognitive in nature.

Perceptual learning of regular auditory patterns is typically rapid, at least in experimental contexts, indicated by faster response times and higher detection sensitivity for recurring than novel patterns, after the same pattern was heard only a few times (Agus et al., 2010; Luo et al., 2013; Kang et al., 2017; Bianco et al., 2020). We would expect sustained neural activity to increase earlier when patterns are recurring compared to novel, if sustained activity reflects perceptual learning of regular patterns. We would further expect that memory for recurring patterns may also affect the magnitude of pattern-related sustained activity. Here, we examine whether the onset time and magnitude of sustained neural activity to regular patterns in tone-pip sequences changes when these pattern sequences reoccur in subsequent trials, compared to when the pattern is heard on only one trial. If sustained activity does depend on experience, it could be further developed as an objective marker of auditory perceptual learning.

Perceptual learning of regular patterns may be implicit: learning may occur when patterns are task irrelevant (Bianco et al., 2020) or when individuals are asleep while sound patterns are presented (Andrillon et al., 2017). Pattern-related increases in sustained neural activity have been observed even when individuals ignore sounds containing patterns (Barascud et al., 2016; Herrmann and Johnsrude, 2018b; Southwell and Chait, 2018), but it is unclear whether changes in sustained activity related to the recurrence of a regular pattern depend on an individual’s attention. Here, we also manipulate attention to examine whether changes in sustained activity related to perceptual learning are observed uniquely when auditory pattern sequences are attended to, or whether they are more automatic and evident even when attention is elsewhere. If attention is not required, this would make sustained activity a robust objective marker to assess perceptual learning and memory formation in populations for which attention cannot be easily manipulated experimentally through instructions.

In a series of five behavioral and EEG experiments, we investigate whether and how sustained activity is affected by the re-occurrence of a regular pattern and whether a person’s attentional state mediates the relationship between pattern recurrence and sustained activity. Perceptual learning is probed by contrasting behavior and sustained neural activity in response to tone-pip patterns that recur throughout the experiment, and similar patterns that are novel on each trial. We expect that the recurrence of regular patterns across trials (compared to novel patterns) provides perceptual benefits, for example, for the detection of patterns and in sensitivity to pattern deviations. Moreover, we expect sustained activity to increase earlier for recurring compared to novel patterns and that the magnitude of sustained activity differs between recurring and novel patterns. This study provides a detailed account of changes in sustained neural activity associated with perceptual learning of auditory patterns.

## General Methods & Materials

### Participants

Participants gave written informed consent prior to the experiment and received course credits or were paid $5 CAD per half-hour for their participation. Participants reported normal hearing abilities. The study was conducted in accordance with the Declaration of Helsinki, the Canadian Tri-Council Policy Statement on Ethical Conduct for Research Involving Humans (TCPS2-2014), and was approved by the local Nonmedical Research Ethics Board of the University of Western Ontario (protocol ID: 106570).

### Stimulation apparatus

Behavioral and EEG recordings were carried out in a sound-attenuating booth. Sounds were presented via Sennheiser (HD 25-SP II) headphones and a Steinberg UR22 (Experiment I) or an RME Fireface 400 (Experiments II-V) external sound card. Stimulation was run using Psychtoolbox in MATLAB (MathWorks Inc.).

### Acoustic stimulation and procedure

At the beginning of an experimental session, a methods-of-limits procedure was used to estimate the participant’s hearing threshold. Details of the procedure are described in detail in previous work (Herrmann and Johnsrude, 2018a). In short, thresholds were obtained by presenting a 1323-Hz sine tone of 12-s duration that either decreased or increased continuously in intensity at 5.8 dB/s. Participants indicated via button press when they could no longer hear the tone (intensity decrease) or when they started to hear the tone (intensity increase). The mean sound intensity at the time of the button press was noted for 6 decreasing tones and 6 increasing tones (decreasing and increasing tones alternated), and these were averaged to determine the individual’s hearing threshold. Acoustic stimuli were presented at 55 dB above the individual’s hearing threshold, that is, at 55 dB sensation level.

Experimental stimuli were 4.8-s long tone sequences that each consisted of 120 tones arranged in twelve sets of ten tones (see also Barascud et al., 2016; Herrmann and Johnsrude, 2018b; Southwell and Chait, 2018). Each set had a duration of 0.4 s (0.04 s individual tone duration; 0.007 s rise time; 0.007 s fall time), with no gap between tones or sets. The frequency of each tone was one of 200 values ranging from 700 to 2500 Hz (logarithmically spaced). Sound sequences were played at 44.1 kHz sampling frequency.

Acoustic stimuli were presented in ‘Novel’ and ‘Recurring’ conditions that were presented with equal probability (50%). For each 4.8-s stimulus, 10 new frequency values were randomly selected for each of the first 4–8 sets (depending on the specific experiment; see below), and then 10 new random frequency values were selected and repeated for the remaining sets (always 12 sets in total), creating a regular pattern. The serial order of tone frequencies was identical in these repeating sets (Figure 1A). Hence, each stimulus started as a sequence of tones with random frequencies, and transitioned part-way through to a regular, repeating pattern of tone frequencies.

**Figure 1:**
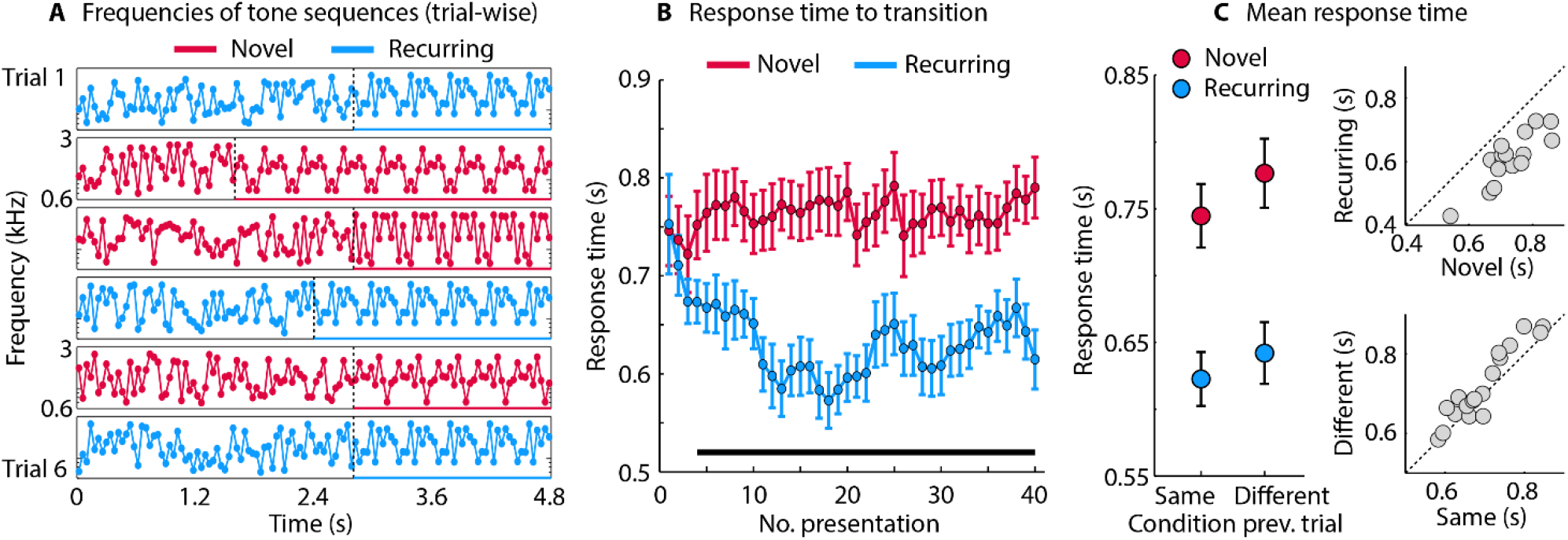
Stimuli and results of Experiment I. **A:** Tone frequencies for six sequential trials in a block of Experiment The black dotted vertical line indicates the onset of a regular pattern and the duration of the regular pattern is indicated by the colored solid line underneath each trial. In the Novel condition, a different regular pattern was presented on each trial. In the Recurring condition, the same regular pattern was present on each trial. The random section of the tone sequences was different from trial to trial for both conditions. **B:** Response times as a function of stimulus presentation (data are smoothed temporally with a 3-point rectangular window). The solid black line indicates a significant difference between novel and recurring regular patterns (p ≤ 0.05; FDR thresholded). **C:** Mean response time across all trials, separately for novel and recurring regular patterns and separately for trials preceded by a trial from the same or different condition.

**Figure 2:**
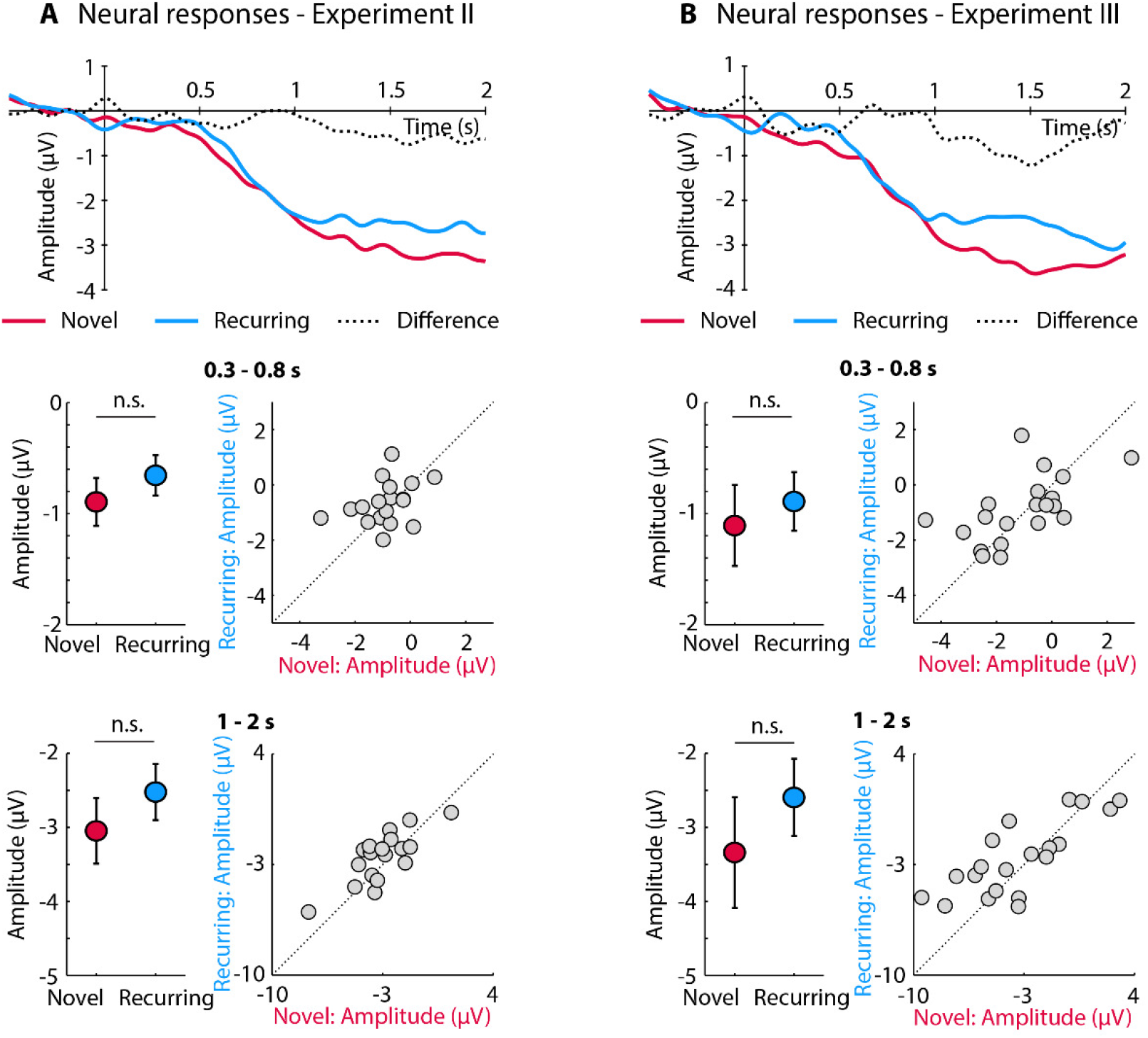
Results for neural responses in Experiment II (A) and III (B). The top row displays neural responses time locked to the onset of the regular pattern in sounds. The two bottom rows show neural responses for Novel and Recurring conditions for the two time windows of interest: 0.3–0.8 s and 1–2 s. Sustained activity did not differ between conditions when data from each experiment were analyzed separately. Pooled data across experiments revealed a significant reduction in sustained activity for Recurring compared to Novel trials in the 1–2 s time window. n.s. –not significant

Critically, in the Novel condition, the set of 10 random frequency values used to create the regular pattern section of the stimulus was selected anew for each trial and for each participant. In the Recurring condition, the regular pattern was identical on successive trials. Hence, trials of both the Novel and Recurring conditions contained a regular pattern, but the pattern was different for each Novel trial and was identical across Recurring trials (although the duration of the regularity changed from trial to trial in both cases, depending on the number of times the pattern set was repeated; between 4 and 8 times).

Participants listened to stimuli in four (Experiment I) or six blocks (Experiments II-V), and could take a break between blocks. In all experiments, randomizations were unique for each participant. The sequence of 10 random frequency values used to create a ‘recurring’ pattern was randomly selected for each participant and for each block, and changed between blocks.

### EEG recordings & analysis

Scalp EEG was recorded from 16 electrodes (Ag/Ag-Cl-electrodes; 10-20 placement) and additionally from the left and right mastoids using a BioSemi EEG system (Amsterdam, The Netherlands). Data were recorded at a sampling rate of 1024 Hz. The online low-pass filter was set at 208 Hz. Electrodes were referenced online to a monopolar reference feedback loop connecting a driven passive sensor and a common-mode-sense (CMS) active sensor, both located posteriorly on the scalp.

MATLAB software (v7.14; MathWorks, Inc.) was used for offline data analysis. Data were filtered with an elliptic filter to suppress line noise at 60 Hz. Data were re-referenced by averaging the two mastoid channels and subtracting the average separately from each of the 16 channels. Data were low-pass filtered at 22 Hz (211 points, Kaiser window) and high-pass filtered at 0.7 Hz (2449 points). Data were divided into epochs ranging from −1 to 5.8 s, time locked to sound onset. Independent components analysis (runica method, Makeig et al., 1996; logistic infomax algorithm, Bell and Sejnowski, 1995; Fieldtrip implementation Oostenveld et al., 2011) was used to identify activity related to blinks and horizontal eye movements. This analysis pipeline was used only for the identification of artifact components.

For the main data analysis, raw data were filtered with the elliptic filter for line noise suppression and with a 7-Hz low-pass filter (501 points, Hann window), re-referenced to the averaged mastoids, before dividing data into epochs ranging from −1 to 5.8 s. High-pass filtering was omitted and data were low-pass filtered at 7 Hz, because sustained activity is a low-frequency response (Barascud et al., 2016; Herrmann and Johnsrude, 2018b; Southwell and Chait, 2018). Blink and eye-movement components from the independent components analysis, identified using the high-pass filtered data, were excluded. Epochs that exceeded a signal change of more than 200 μV for any electrode were excluded from analyses.

In order to analyze differences in sustained activity between the Novel and Recurring conditions following the transition from the random section to the regular section of the stimulus, we extracted data epochs ranging from −0.5 to 2 s time-locked to the transition onset within the sound. Single-trial time courses were averaged separately for Novel and for Recurring trials. Mean time courses were baseline-corrected by subtracting the mean signal in the −0.5 to 0 s time window from the signal at each time point of the epoch (separately for each channel). Data were averaged across a fronto-central electrode cluster (F4, Fz, F3, C4, Cz, C3) that we know from previous work to be sensitive to regularity-related sustained activity (Herrmann and Johnsrude, 2018b).

Data analysis focused on two time windows. An early time window, ranging from 0.3 to 0.8 s post-transition onset, was used to determine whether regularity-related sustained activity is exhibited earlier for the Recurring than for the Novel condition. This time window was chosen because it takes about 1.5 sets (0.6 s) to elicit regularity-related sustained activity after transitioning from a random to a (novel) regular tone sequence (Barascud et al., 2016). Neural-response amplitudes within the 0.3–0.8-s time window were averaged. Latency analysis was not possible, because of the slow and sustained nature of response, but any difference in response latency between two time courses will be reflected in amplitude changes for a fixed analysis time window. The second time window, ranging from 1 to 2 s post-transition onset, was used to determine whether the magnitude of regularity-related sustained activity differs between the Recurring and the Novel conditions. Neural-response amplitudes within the 1–2-s time window were averaged. Condition effects (Novel versus Recurring) in these two time windows have opposing polarities (Figure 4). To avoid averaging data in the transition period between the two time windows, we did not analyze neural activity from 0.8 to 1 s.

**Figure 3:**
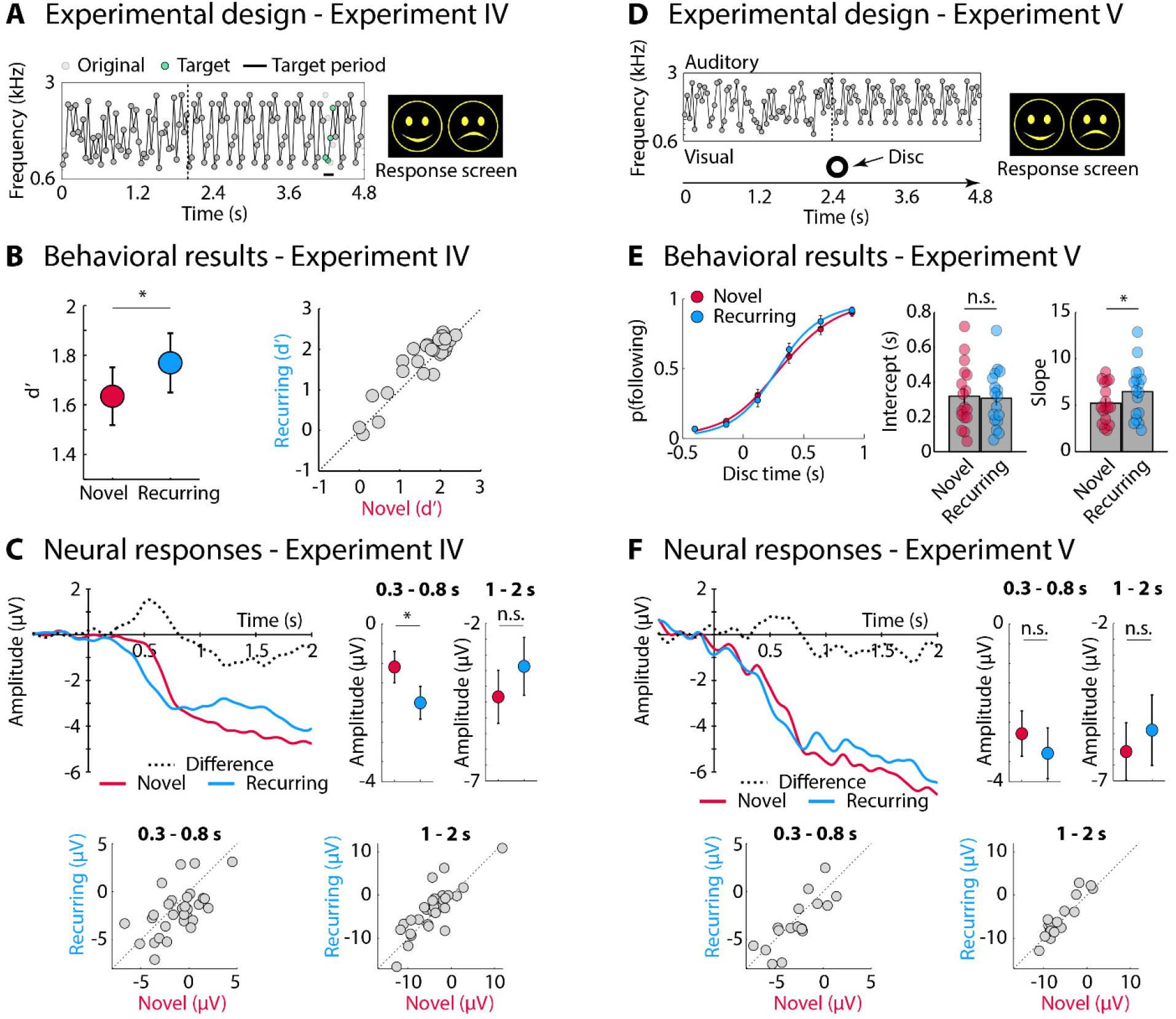
Experimental design and results for Experiment IV and V. **A:** Experimental design for Experiment IV. A deviation from the regular pattern was inserted into half of the trials by replacing several tone frequencies (median: 4 frequencies), in the 11th set of the sound. The example shows the frequencies for a sound that included a deviant; deviant is marked by the colored dots and the black horizontal line. After sound presentation, participants pressed the key for the happy smiley-face icon (deviation was present) or the key for the sad smiley-face icon (deviation was absent). **B:** Behavioral results indicated that sensitivity was higher for trials in the Recurring compared to Novel condition. **C:** Neural responses: time courses and mean activity. For the 0.3-0. 8 s time window, sustained activity increased earlier (i.e., measured as a more negative amplitude) in the Recurring than Novel condition. **D:** Experimental design for Experiment V. On each trial, a visual disc was presented at one of six times relative to the transition to a regular pattern within the sound. After sound presentation, participants pressed the key for the happy smiley-face icon (disc appeared after transition to regular pattern) or the key for the sad smiley-face icon (disc appeared before transition to regular pattern). **E:** Behavioral results indicated a steeper slope for the Recurring compared to the Novel condition. **F:** Neural responses: time courses and mean activity for the two time windows of interest (0.3-0.8 s and 1-2 s; there was no significant difference). *p ≤ 0.05, n.s. - not significant

**Figure 4:**
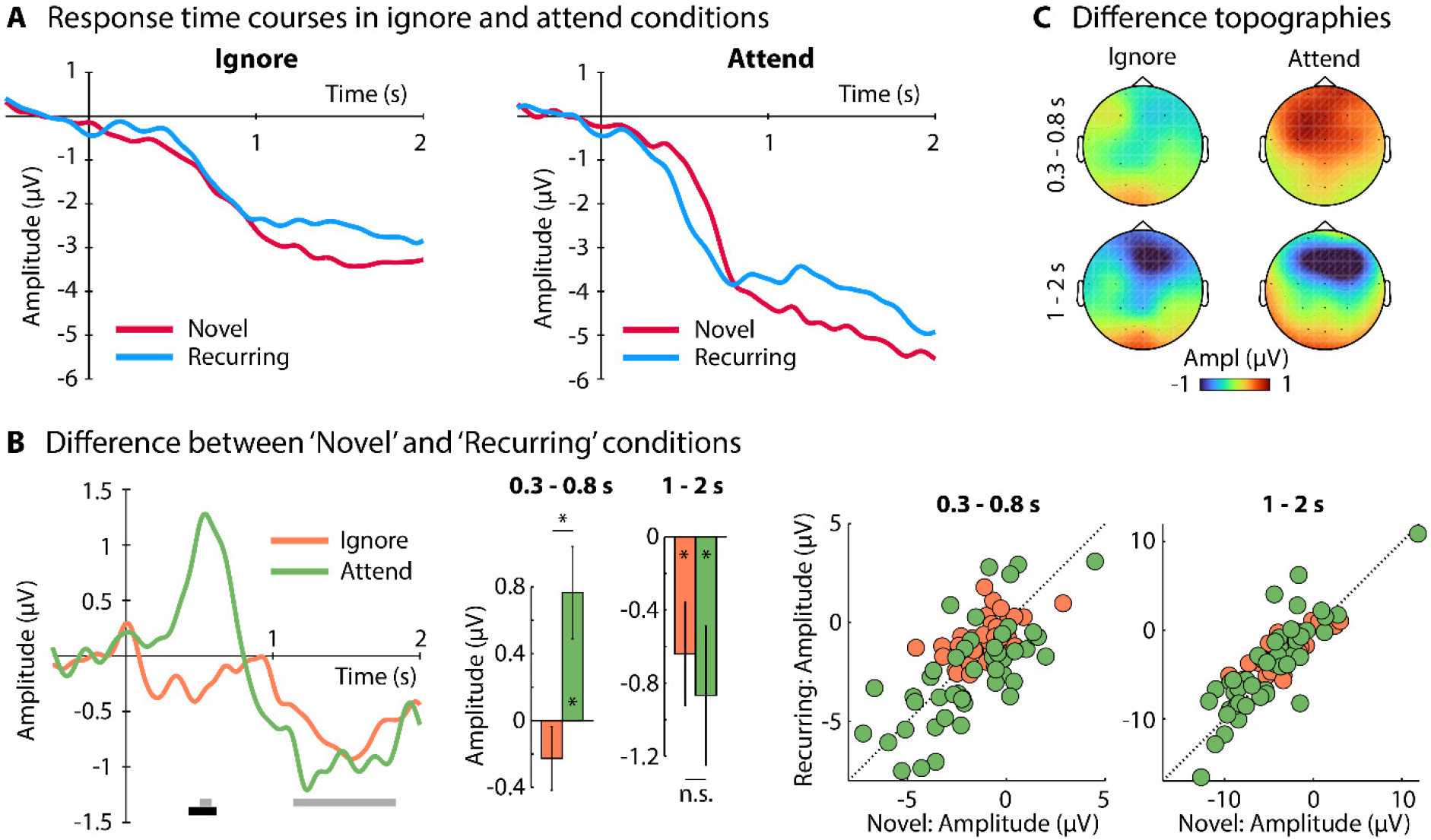
Effects of attention on regularity-related sustained activity in Novel and Recurring conditions. **A:** Time courses (collapsed across the two experiments in each case: Ignore: Experiments II and III; Attend: Experiments IV and V). **B:** Differences between Novel and Recurring conditions: Difference time courses for ignore and attend conditions are shown on the left. The gray line below the time courses reflects time points at which the main effect of Condition (Recurring vs. Novel) was significant (difference from zero; FDR-thresholded). The black line reflects time points at which the Condition × Attention interaction was significant (difference between ignore and attended; FDR-thresholded). Bar graphs for time windows of interest are shown (error bar reflects the standard error of the mean). The asterisk within bars indicates a significant difference between Novel and Recurring conditions. Individual data points are displayed on the right (each dot reflecting data from one participant). For the 0.3-0.8 s time window, sustained activity was smaller (i.e., more positive) for the Novel than Recurring conditions when participants attended to sounds, but not when they ignored sounds (interaction p = 0.006). For the 1-2 s time window, sustained activity was reduced for the Recurring compared to the Novel condition, both when participants attended to, and ignored the sounds. **C:** Topographical distributions of the amplitude difference between Novel and Recurring conditions. *p ≤ 0.05, n.s. - not significant

### Experimental design and statistical analysis

A paired samples t-test was used to compare amplitudes between the Novel and the Recurring conditions, separately for the 0.3-0.8 s and the 1-2 s time windows. ANOVAs were used to investigate effects of conditions across experiments (see below). Effect sizes are provided as partial η^2^ for ANOVAs and as r_e_ (r_equivalent_) for t-tests (Rosenthal and Rubin, 2003). r_e_ is equivalent to the square root of partial η^2^ for ANOVAs. We further provide Inclusion Bayes Factors (BF_incl_) using the “matched models” procedure in JASP software (van den Bergh et al., 2019; JASP, 2020). This study was not pre-registered. Data in BIDS format (Pernet et al., 2019) are available at https://osf.io/9fmz5/.

## Experiment I: Faster detection for recurring than novel regular patterns

Experiment I aimed to replicate previous behavioral work that demonstrates improved detection of a recurring compared to a novel regular pattern (Agus et al., 2010; Bianco et al., 2020) so that we can be sufficiently confident about the sensitivity of our experimental paradigm for characterization of behavioral benefits and neural signatures of perceptual learning.

### Methods and materials

#### Participants

Nineteen participants took part in Experiment I (median age: 20 years; range: 19–22 years; 6 female). Data from one additional participant were excluded because no response was made in more than 25% of trials.

#### Stimuli and procedure

Participants listened to the 4.8-s tone sequences in four blocks. Each block included 80 sequences, one in each of 80 trials: 40 Novel (different regular pattern on each trial) and 40 Recurring (same regular pattern on each trial). Within each trial, tone sequences transitioned from the random section to the regular pattern at the beginning of the 5th, 6th, 7th, 8th, or 9th, set of 10 tones (counterbalanced across conditions), corresponding to 1.6, 2, 2.4, 2.8, or 3.2 s after onset, respectively. Trials within each block were randomized such that a maximum of three trials from one condition (Novel or Recurring) could occur in a row. Participants were instructed to respond as quickly as possible via key press when they detected the regular pattern. Only responses made within 2 s after pattern onset were considered. Response times were estimated relative to the onset of the first tone of the regular pattern (i.e., 1.6, 2, 2.4, 2.8, or 3.2 s). The intertrial interval was 2 s.

To measure simple change-detection response time, participants heard twenty 4.8-s sounds, in which constituent tones were fixed at one of two frequencies in a single control block. The tones of the first 5, 6, 8, or 9 sets were fixed at 1323 Hz; tones in the remaining sets were all fixed at 1600 Hz. Participants indicated via button press, as quickly as possible, when the switch from one fixed frequency to another happened. Response times were averaged across control trials to estimate the response time to a non-demanding stimulus change, so as to estimate the time it takes for a change to reach awareness and to be translated into motor output (Barascud et al., 2016).

The mean response time from the control block was subtracted from the response time of each trial in the Novel and Recurring conditions in order to normalize response times for the time of awareness and motor processes. For each condition, response times were averaged across the four stimulation blocks, separately for each of the 40 trials per condition presented within a block. This led to two 40-trial response time courses as a function of stimulus presentation. Response times were smoothed using a 3-point rectangular window to increase the signal-to-noise ratio of the mean response time at each point of the time course. A paired t-test was used to compare response times between the Novel and the Recurring conditions at each of the 40 trial positions. False discovery rate was used to account for multiple comparisons (Benjamini and Hochberg, 1995; Genovese et al., 2002).

Considering that a maximum of three trials of the same condition could occur in a row, we also investigated whether any response-time benefit for recurring over novel regular patterns may be driven by trials that are preceded by the same condition or whether the response-time benefit is also presented for trials preceded by a different condition. Response times were analyzed using a repeated measures ANOVA with factors Condition (Novel, Recurring) and Previous Condition (Same, Different).

### Results

Response times associated with the stimulus presentation order are depicted in Figure 1B. At the very beginning of a stimulation block, response times did not differ between Novel and Recurring trials. By the time participants had heard 4-5 occurrences of the same regular pattern (in the Recurring condition), they responded significantly faster compared to novel regular patterns (Figure 1B).

The repeated-measures ANOVA revealed faster response times for recurring compared to novel patterns (effect of Condition: F_1,18_ = 162.958, p < 1^e-8^, η^2^_p_ = 0.902, BF_incl_ = 4.86^e+17^) and faster response times when the previous trial was the same compared to the different condition (effect of Previous Condition: F_1,18_ = 12.581, p = 0.002, η^2^_p_ = 0.411, BF_incl_ = 9.807), but no interaction (F_1,18_ = 0.655, p = 0.429, η^2^_p_ = 0.035, BF_incl_ = 0383; Figure 1C). The data suggest that the response-time benefit for recurring regular patterns is not due entirely to the same condition on a previous trial, but that individuals detect the onset of recurring patterns faster even when novel patterns are presented in-between successive Recurring trials.

The slower response times for trials preceded by a different condition may indicate a violation from an expected pattern. Since this did not depend on whether the preceding pattern was novel or recurring, we conclude that individuals have formed some persistent representation of the previous trial, even if it was novel.

The results of Experiment I are in line with previous work and show that individuals learn regular tone patterns over a few trials (Agus et al., 2010; Kang et al., 2017; Bianco et al., 2020), such that they are faster in detecting the pattern when they encounter it again (Bianco et al., 2020). Participants detected a recurring regular pattern about 0.12 s earlier than a novel regular pattern.

In the subsequent EEG experiments (II-V), we explore whether this perceptual benefit also manifests as earlier sustained activity and/or a change in magnitude of sustained activity in response to recurring compared to novel regular patterns, and whether attention to sounds is needed for changes in sustained activity related to perceptual learning.

## Experiment II and III: Sustained activity during passive listening to novel and recurring patterns

Regularity-related sustained activity is typically investigated in passive-listening paradigms or under visual distraction (Barascud et al., 2016; Teki et al., 2016; Herrmann and Johnsrude, 2018b; Southwell and Chait, 2018). Experiment II and III of the current study thus investigate whether or not sustained activity is different for recurring compared to novel patterns under passive listening conditions. Novel and Recurring trials were randomly interspersed within blocks in Experiment II and blocked in Experiment III.

### Methods and Materials

#### Participants

Eighteen individuals took part in Experiment II (median age: 21 years; range: 18–29 years; 13 female) and twenty different individuals participated in Experiment III (median age: 18 years; range: 18–23 years; 17 female). None of the participants who took part in Experiment II and III took part in Experiment I. Data from four additional participants were recorded for Experiment II, but due to technical problems no triggers were recorded, and the data could not be analyzed.

#### Stimuli and procedure

In Experiments II and III, EEG was recorded while participants listened passively to auditory stimuli and watched a muted, subtitled movie of their choice on a portable DVD player.

The stimulation procedure for Experiment II was identical to that in Experiment I, with a few exceptions. In each of six blocks, participants listened to 102 trials, 51 in each of the Novel and Recurring conditions. Sounds transitioned from the random section to the regular pattern at set 6, 7, or 8 (with equal probability; that is 17 trials each per block and condition). Participants thus listened to a total of 306 trials per Novel and Recurring condition. The inter-stimulus interval was 2 s.

In Experiment III, trials from the Novel and Recurring conditions were blocked in order to investigate whether this would increase the effects of pattern re-occurrence. In each of six blocks, participants listened to 72 trials. In three blocks, 36 Novel trials preceded 36 Recurring trials, and this condition order was reversed in the other three blocks. Blocks with Novel versus Recurring trials being presented first alternated. The order of the six blocks was counterbalanced across participants. Tone sequences of the Novel and Recurring conditions transitioned to a regular pattern at set 5, 6, or 7 (with equal probability). Half of the sounds also contained a pattern deviation that was generated by replacing the frequency of 4 out of the 120 tones making up a sound sequence (in set 11; tones 105 to 108) with 4 different randomly selected frequencies. The same procedure was used in Experiment IV, where this pattern deviation served as a detection target (Figure 3A).

### Results

In Experiment II, neural responses did not differ between the Novel and the Recurring condition for the 0.3–0.8 s time window (t_17_ = 1.043, p = 0.312, r_e_ = 0.245, BF_incl_ = 0.505) nor for the 1–2 s time window (t_17_ = 1.906, p = 0.074, r_e_ = 0.420, BF_incl_ = 1.145; Figure 2A). The results were qualitatively similar for Experiment III, in which trials were blocked by condition: neural responses did not differ between the Novel and the Recurring conditions for the 0.3–0.8 s time window (t_19_ = 0.715, p = 0.483, r_e_ = 0.162, BF_incl_ = 0.372) nor for the 1–2 s time window (t_19_ = 1.568, p = 0.133, r_e_ = 0.339, BF_incl_ = 0.747; Figure 2B).

The top part of Figure 2A and 2B shows that, despite the absence of a statistically significant effect in the 1–2 s time window, sustained activity was less negative (i.e., reduced) in the Recurring compared to the Novel condition in both experiments. This suggest that a true effect may be present, but that the effect is small in each individual experiment and requires greater statistical power. In order to increase statistical power, data from both experiments were submitted to an explorative ANOVA with the within-subjects factor Condition (Novel, Recurring) and the between-subjects factor Experiment (Experiment II, III). For the 1–2s time window, this analysis revealed a significant reduction in sustained activity for the Recurring compared to the Novel condition (F_1,36_ = 5.049, p = 0.031, η^2^_p_ = 0.123, BF_incl_ = 1.988). The effect of Experiment (F_1,36_ = 0.061, p = 0.806, η^2^_p_ = 0.002, BF_incl_ = 0.478) and the Condition × Experiment interaction (F_1,36_ = 0.151, p = 0.700, η^2^_p_ = 0.004, BF_incl_ = 0.354) were not significant. No significant main or interaction effects were observed for the 0.3–0.8 s time window (all F < 1.4, p > 0.2, η^2^_p_ < 0.04, BF_incl_ < 0.5). Hence, we observe a small reduction in sustained activity in the Recurring compared to Novel condition.

In sum, the results of Experiment II and III do not reveal earlier sustained activity for the Recurring compared to the Novel condition (i.e., no effects for the 0.3–0.8 s time window, Figure 2A,B top). The analyses, however, revealed a small reduction in sustained activity for recurring compared to novel regular patterns in the 1–2 s time window (data pooled across experiments). This reduction in the magnitude of sustained activity may not be directly related to perceptual learning (which was predicted to lead to earlier sustained activity), but may relate to processes enabled by learning.

Previous work has demonstrated an increase in the magnitude of sustained activity for sounds with a regular pattern compared to sounds without a regular pattern under passive or distracted listening conditions (Barascud et al., 2016; Herrmann and Johnsrude, 2018b; Southwell and Chait, 2018), but this work did not investigate perceptual learning of regular patterns over multiple trials. Attention to the patterned sounds may be needed for individuals to learn patterns (Huyck and Johnsrude, 2012), such that the associated sustained activity occurs earlier compared to novel patterns. Accordingly, in Experiments IV and V, we required individuals to attend to the sounds.

## Experiment IV and V: Earlier sustained activity for recurring than novel patterns under attention

Experiments IV and V were designed to investigate whether sustained activity increases earlier for recurring compared to novel patterns when participants attend to the sounds. We utilized two different active tasks and examined changes in behavior in these tasks for recurring compared to novel patterns. In Experiment IV, participants had to detect a near-threshold deviation from a regular pattern. We expected that deviation detection sensitivity may be higher for recurring than for novel patterns, if learning of the recurring pattern strengthens its representation. In Experiment V, participants judged, for each sound, whether a visual disc, presented at one of several fixed times relative to pattern onset, precedes or follows the onset of the pattern. We expected that learning of the recurring pattern would lead to a greater proportion of ‘follow’ responses for the visual stimulus, since Experiment I (Figure 1) and previous observations (Bianco et al., 2020) demonstrate that individuals detect a recurring pattern faster than a novel pattern. In Experiment IV and V, we also expected to see that sustained activity increases earlier for recurring compared to novel patterns. The procedures of Experiment I could not be implemented for an active EEG task, because anticipation of a motor response increases sustained activity (Jahanshahi and Hallett, 2003; Lang, 2003) that may confound pattern-related sustained activity, and the motor response would occur at the time when pattern-related sustained activity emerges.

### Methods and Materials

#### Participants

Thirty-two individuals took part in Experiment IV (median age: 18 years; range: 18–29 years; 16 female) and seventeen different individuals participated in Experiment V (median age: 18 years; range: 17–21 years; 13 female). None of the participants took part in any other experiment. Data from one additional participant for Experiment IV were recorded, but the corresponding log files for half of the experimental blocks were not stored due to technical problems. Data for this participant were thus not analyzed. Note that COVID-19 related restrictions to data recording contributed to differences in the number of participants across experiments. We had originally aimed for about 30 participants for Experiment V as well.

#### Stimuli and procedure

Acoustic stimulation in Experiment IV was similar to Experiment III: Novel and Recurring trials were blocked, with half of them including a frequency deviation in the 11th set. In this experiment, participants were asked to indicate the detection of the frequency deviation with a keypress. Since preparation of a motor response could influence low-frequency sustained activity (Jahanshahi and Hallett, 2003; Lang, 2003), the behavioral response was delayed, and cued visually 0.01 s after sound offset. The visual cue consisted of a happy and a sad smiley-face icon presented side by side (Figure 3A). The happy smiley-face icon indicated the button for ‘deviation present’, whereas the sad smiley-face icon indicated the button for ‘deviation absent’. The position of the happy and sad smiley-face icons (left vs. right) was random and participants did therefore not know which button to press prior to the visual response screen, and could thus not prepare any specific motor response (Herrmann et al., 2011a; Herrmann et al., 2011b). The next trial started 2.2 s after they had made a response. Participants performed 2-4 short training blocks to familiarize them with the task and to titrate the number of altered tones that made up the deviant so that participants would reach an ~80% detection rate (a median of 4 tones were used for the deviation).

The acoustic stimulation in Experiment V was similar to that in Experiments III and IV: Sounds had a duration of 4.8 s, each containing a regular pattern starting at set 6, 7, or 8 (with equal probability). Either a new pattern (Novel) or same pattern (Recurring) was presented on each trial, and trials for each condition (Novel, Recurring) were clustered within a stimulation block. On each trial, a disc (white on black background) was presented visually for 0.15 s at one of six times relative to the transition at which the sound changed to a regular pattern: −0.4, −0.14, 0.12, 0.38, 0.64, 0.9 s. We included more time points following than preceding the transition to a regular pattern, because the detection of a pattern requires the accumulation of evidence over a few hundred milliseconds (Barascud et al., 2016). Participants were asked to indicate whether the disc appeared after the onset of the regular pattern (Vroomen and Keetels, 2010; Patel and Chait, 2011). As in Experiment IV, the response was cued visually 0.01 s after sound offset, to avoid motor preparation. If the disk appeared after the sound transition, they pressed the button indicated by the happy smiley-face icon (“disc following pattern onset”); otherwise, they pressed the button indicated by the sad smiley-face icon (Figure 3A). The position of the two icons (left vs. right) was random. The next trial started 2.2 s after the participant had made a response. Participants performed 1-2 short training blocks to familiarize them with the task.

#### Behavioral data analysis

For Experiment IV, perceptual sensitivity (d’) was calculated for Novel and Recurring conditions. A hit was defined as a correct ‘deviation present’ response. A false alarm was defined as an incorrect ‘deviation present’ response (i.e., when no deviation was present). A repeated-measures ANOVA with factors transition time (set 5, 6, or 7) and condition (Novel, Recurring) was calculated.

For Experiment V, the proportion of ‘following’ (disc after pattern onset) responses was calculated for each of the six disc times (i.e., −0.4, −0.14, 0.12, 0.38, 0.64, 0.9 s) and two sound conditions (Novel, Recurring). For each participant, a logistic function was fit to the proportion data as a function of disc time relative to pattern onset, with intercept and slope as free parameters. Differences between Novel and Recurring conditions were tested separately for the intercept and slope using paired sample t-tests.

### Results

For Experiment IV, participants were more sensitive to deviations in the regular pattern when the transition to a regular pattern started earlier compared to later in the sound (F_2,62_ = 5.525, p = 0.011, η^2^_p_ = 0.151, ε = 0.793, BF_incl_ = 7.939), suggesting that the regular pattern is better represented when a higher number of regular sets occurred prior to the deviation. Critically, participants were more sensitive to deviations in the regular pattern for recurring compared to novel patterns (F_1,31_ = 8.308, p = 0.007, η^2^_p_ = 0.211, BF_incl_ = 45.217; Figure 3B; the interaction was not significant, p > 0.4). This suggests that the re-occurrence of a regular pattern across trials also enhanced the representation of the regular pattern.

Analysis of neural responses revealed earlier sustained activity (measured as amplitude in the 0.3–0.8 s time window) for the Recurring compared to the Novel condition (t_31_ = 2.458, p = 0.0198, r_e_ = 0.404, BF_incl_ = 2.607; Figure 3C). For the 1–2 s time window, regularity-related sustained activity was reduced (less negative) in the Recurring compared to the Novel condition, but this difference was not significant (t_31_ = 1.810, p = 0.080, r_e_ = 0.309, BF_incl_ = 0.931).

In Experiment V, the estimated intercept of the logistic function between the Recurring and the Novel condition did not differ (t_16_ = 0.496, p = 0.627, r_e_ = 0.123, BF_incl_ = 0.357). However, the slope of the logistic function fit was larger (i.e., steeper) for the Recurring compared to the Novel condition (t_16_ = 2.464, p = 0.0255, r_e_ = 0.525, BF_incl_ = 2.412; Figure 3E), indicating that the timing of the visual disc relative to pattern onset could be estimated more accurately on Recurring trials. This may indicate a shorter audio-visual temporal integration window for the former relative to the latter (Vroomen and Keetels, 2010).

Sustained neural activity did not differ between Recurring and Novel conditions for the 0.3–0.8 s time window (t_16_ = 1.286, p = 0.217, r_e_ = 0.306, BF_incl_ = 0.583) or for the 1–2 s time window (t_16_ = 1.528, p = 0.146, r_e_ = 0.357, BF_incl_ = 0.758; Figure 3F).

As for the analysis of sustained activity for the passive-listening Experiments II and III, Figure 3 shows that, for both Experiments IV and V, sustained activity was less negative (i.e., reduced) in the 1–2 s time window. Although this was not significant using two-tailed tests in either experiment, it appeared consistent across experiments, and so, as for Experiments II and III, we pooled the data from Experiment IV and V and conducted an explorative ANOVA with the within-subjects factor Condition (Novel, Recurring) and the between-subjects factor Experiment (Experiment IV, V).

For the 1–2 s time window, as in Experiments II/III, sustained activity was smaller (i.e., less negative) in the Recurring compared to the Novel condition (F_1,47_ = 4.199, p = 0.046, n^2^_p_ = 0.082, BF_incl_ = 1.897), but there was no effect of Experiment (F_1,47_ = 1.906, p = 0.174, n^2^_p_ = 0.039, BF_incl_ = 0.801) nor a Condition × Experiment interaction (F_1,47_ = 0.128, p = 0.723, n^2^_p_ = 0.003, BF_incl_ = 0.359).

For the 0.3–0.8 s time window, sustained activity was overall more negative in Experiment V than Experiment IV (effect of Experiment: F_1,47_ = 5.209, p = 0.027, n^2^_p_ = 0.100, BF_incl_ = 2.979). Critically, sustained activity was more negative – indicating shorter latency – in the Recurring compared to the Novel condition (F_1,47_ = 5.855, p = 0.019, n^2^_p_ = 0.111, BF_incl_ = 5.793), but there was no Condition × Experiment interaction (F_1,47_ = 0.497, p = 0.484, n^2^_p_ = 0.010, BF_incl_ = 0.297). The earlier increase in sustained activity due to pattern recurrence is consistent with the faster detection of recurring compared to novel regular patterns in Experiment I, which indexes perceptual learning (Figure 1).

## Examining whether attention affects differences between recurring and novel patterns

The results from the previous sections demonstrate reduced regularity-related sustained activity for the Recurring compared to the Novel condition for the 1–2 s time window regardless of whether sounds are ignored (Experiments II and III) or attended (Experiments IV and V). This effect appears to be small, but it was reliably present in the averaged time courses in each of the four EEG experiments (Figures 2A/B and 3C/F). An earlier increase in regularity-related sustained activity (measured as amplitude difference in the 0.3–0.8 s time window) for the Recurring compared to the Novel condition was observed only when participants attended to the sounds (Experiments IV/V, but not Experiments II/III). In order to test directly whether sustained activity manifests earlier for recurring compared to novel patterns when stimuli are attended, we compared these two sets of experiments explicitly using ANOVAs with the within-subjects factor Condition (Novel vs. Recurring) and the between-subjects factor Attention (Ignore [Experiments II and III] vs. Attend [Experiments IV and V]), separately for the 0.3-0.8 s and 1-2 s time windows.

## Sustained activity increases earlier for recurring than novel patterns only when individuals attend to sounds

For the 0.3–0.8 s time window, the Condition × Attention interaction was significant (F_1,85_ = 7.835, p = 0.006, η^2^_p_ = 0.084, BF_incl_ = 6.681): Recurring compared to novel patterns led to earlier sustained activity (measured as amplitude in the 0.3–0.8 s time window) only when participants attended to the sounds (Figure 4). The main effect of Attention indicated larger sustained activity when participants attended to the sounds compared to when they ignored them (F_1,85_ = 8.77, p = 0.004, η^2^_p_ = 0.094, BF_incl_ = 8.938). There was no main effect of Condition (F_1,84_ = 0.001, p = 0.971, η^2^_p_ < 0.001, BF_incl_ = 0.720). In order to ensure that the Condition × Attention interaction was not driven by differences in stimulation protocols between experiments, we conducted the same ANOVA using only data from Experiment III (ignore) and Experiment IV (attend), for which the sound stimulation protocol was the same. For this analysis, we again observed a Condition × Attention interaction (F_1,50_ = 4.571, p = 0.037, η^2^_p_ = 0.084, BF_incl_ = 1.987), demonstrating that sustained activity increased earlier for recurring compared to novel patterns only when participants attended to the sounds.

For the 1–2 s time window, the ANOVA revealed reduced sustained activity for the Recurring compared to the Novel condition (main effect of Condition: F_1,85_ = 9.247, p = 0.003, η^2^_p_ = 0.098, BF_incl_ = 11.545), but attention did not appear to modulate the condition effect (Condition × Attention: F_1,85_ = 0.210, p = 0.648, η^2^_p_ = 0.002, BF_incl_ = 0.252; Figure 4). The Condition × Attention was also not significant when only data from Experiments III and IV were used (F_1,50_ = 0.083, p = 0.775, η^2^_p_ = 0.002, BF_incl_ = 0.295; main effect of Condition: F_1,50_ = 4.887, p = 0.032, η^2^_p_ = 0.089, BF_incl_ = 2.147). Critically, the significant effect of Condition when the data from all four EEG experiments are included in the analysis as well as the presence of the effect in the averaged time courses in each of the four experiments (Figures 2 and 3) highlights that the effect is small, but reliable.

To further indicate that the effects reported here are robust, we calculated the effect of Condition and the Condition × Attention interaction for each time point using False Discovery Rate multiple-comparisons correction (Benjamini and Hochberg, 1995; Genovese et al., 2002; lines below time courses in Figure 4B). These analyses mirrored the analyses for the two time windows, showing a Condition × Attention interaction from about 0.4 to 0.6 s after pattern onset and a main effect of Condition from about 1.1 to 1.8 s (Figure 4B).

Our analyses have focused on a fronto-central electrode cluster. An exploratory analysis tested whether the relevant effects involving Condition (Novel vs. Recurring) differed between frontal and central electrodes. For the 0.3–0.8 s time window, the Condition × Attention interaction did not differ between frontal and central electrodes (F_1,85_ = 0.08, p = 0.778, η^2^_p_ = 0.001). For the 1–2 s time window, in contrast, the effect of Condition was larger for frontal compared to central electrodes (F_1,85_ = 20.357, p = 0.00002, η^2^_p_ = 0.193; effect of Condition for frontal electrodes: F_1,85_ = 16.703, p = 0.000099, η^2^_p_ = 0.164, BF_incl_ = 308.864; effect of Condition for central electrodes: F_1,85_ = 3.264, p = 0.074, η^2^_p_ = 0.037, BF_incl_ = 0.709; see also Figure 4C).

## Earlier latency of sustained activity mirrors behavioral pattern of rapid perceptual learning

To investigate the evolution of the difference between Recurring and Novel trials in sustained activity over the course of a block, trials of each condition (across experiments) were separated into two groups according to whether they occurred early (first half of the trials) or late (second half of trials) within a stimulation block, and then averaged. An ANOVA was calculated using the within-subjects factors Condition (Novel vs. Recurring) and Time (Early vs. Late), separately for the two time windows of interest.

In the 0.3–0.8 s time window, the difference between the Novel and the Recurring conditions was significant for both early and late trials (main effect of Condition: F_1,48_ = 8.443, p = 0.006, η^2^_p_ = 0.150, BF_incl_ = 29.828; early: t_48_ = 2.056, p = 0.045, r_e_ = 0.285, BF_incl_ = 1.282; late: t_48_ = 2.603, p = 0.012, r_e_ = 0.352, BF_incl_ = 3.596; Figure 5A), with no effect of Time or interaction (p > 0.05). The observation that sustained activity increased earlier for the Recurring compared to the Novel condition, and that this increase is apparent in the first half of the presented trials, is consistent with the behavioral results of Experiment I (Figure 1), indicating rapid perceptual learning.

**Figure 5:**
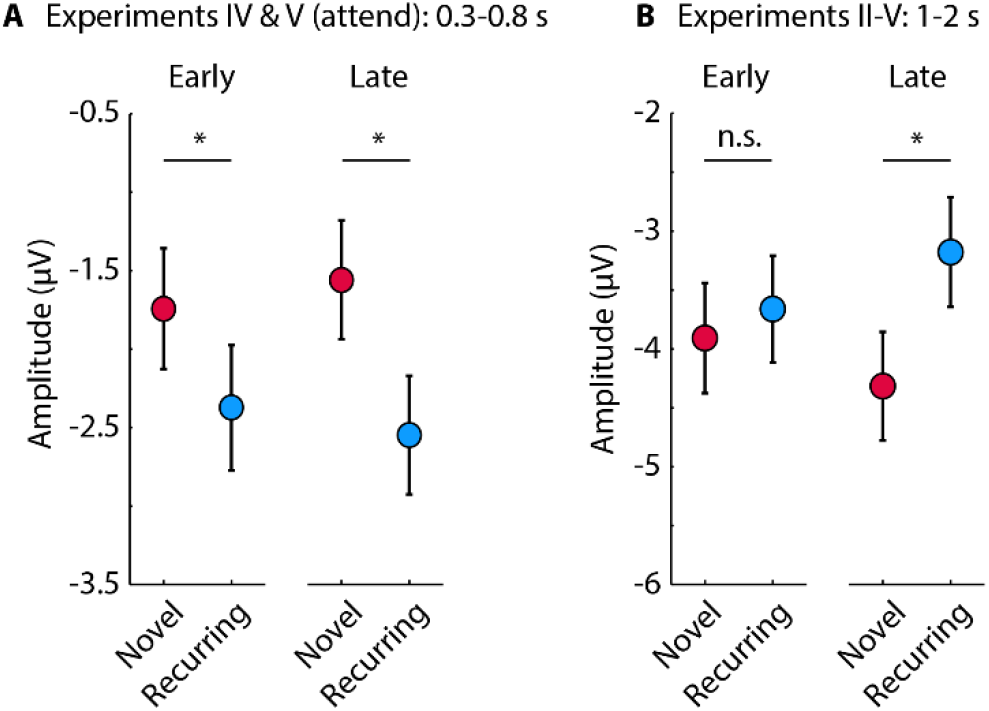
Comparison of sustained activity for Novel and Recurring conditions for early and late trials. **A:** Mean responses in the 0.3-0.8 s time window (Attend condition) were significantly increased (more negative) for the Recurring compared to the Novel condition for early as well as for later trials within a stimulation block. **B:** Mean responses in the 1-2 s time window (Attend and Ignore conditions) were significantly reduced (less negative) for the Recurring compared to the Novel condition for late, but not for early trials within a stimulation block. Error bars reflect the standard error of the mean. *p ≤ 0.05, n.s. - not significant

In the 1–2 s time window, the Condition × Time interaction was significant (F_1,86_ = 4.233, p = 0.043, η^2^_p_ = 0.047, BF_incl_ = 0.786; Figure 5B), with only late trials exhibiting a difference in sustained activity between Recurring and Novel trials (t_86_ = 3.158, p = 0.002, r_e_ = 0.322, BF_incl_ = 12.923). No difference was evident for early trials (t_86_ = 0.822, p = 0.413, r_e_ = 0.088, BF_incl_ = 0.294; Figure 5B). Note that the results are similar if only frontal electrodes are used. That sustained activity is reduced for the Recurring compared to the Novel condition only for late trials appears to differ from the behavioral results of Experiment I where reaction time differences were apparent already after a few trials (Figure 1). This may suggest that the reduced magnitude of sustained activity due to pattern re-occurrence over trials may not be directly related to perceptual learning, but processes that are enabled by learning.

In sum, these data suggest a functional difference between the onset of sustained activity which is sensitive to rapid perceptual learning when a person attend to sounds, and its magnitude, which seems to reflect processes that result from learning independent of a person’s attentional state.

## General discussion

Here we investigated whether and how perceptual learning of regular patterns in sounds changes sustained neural activity, which is a well-established index of pattern processing (Gutschalk et al., 2002; Barascud et al., 2016; Southwell et al., 2017; Herrmann and Johnsrude, 2018b).

### Behavioral evidence of pattern learning

We observed several behavioral benefits associated with the recurrence of a regular pattern that suggest perceptual learning. Detection of regular patterns was faster (Experiment I; Figure 1), sensitivity to pattern deviations higher (Experiment IV; Figure 3), and judgements about the temporal order of pattern onset relative to a visual stimulus more accurate (Experiment V; Figure 3) when participants listened to recurring compared to novel patterns. Perceptual benefits emerged rapidly after only a few recurrences of a regular pattern (Experiment I, Figure 1), which is consistent with previous work (Hawkey et al., 2004; Agus et al., 2010; Agus and Pressnitzer, 2013; Luo et al., 2013; Kang et al., 2017; Bianco et al., 2020). Moreover, faster response times for recurring than novel patterns in Experiment I were observed despite novel patterns being interspersed among recurring patterns, indicating longer-term memory of specific patterns, despite interference by other patterns with an identical temporal structure (cf. Agus et al., 2010; Viswanathan et al., 2016; Bianco et al., 2020).

Participants detected recurring regular patterns faster than novel patterns in Experiment I (Figure 1). We therefore expected when participants had to judge whether a visual disc preceded or followed the onset of the pattern (Experiment V) that they would indicate an earlier disc time for onsets of recurring patterns compared to novel patterns. However, this was not observed (Figure 3E). Earlier detection of a pattern onset was advantageous for performance in Experiment I, whereas Experiment V may not have provided such an obvious advantage because participants focused on audio-visual integration of the auditory pattern and the visual disc. Audio-visual integration may be a more complex process compared to auditory pattern detection alone, potentially decreasing sensitivity to changes in perceived timing. Interestingly, we observed that temporal order sensitivity (i.e., the slope of the psychometric function) – increased for recurring compared to novel patterns (Figure 3E), which may indicate a narrower audio-visual temporal integration window for the former relative to the latter (Vroomen and Keetels, 2010).

### Early increase in sustained activity mirrors rapid perceptual learning

Sustained neural activity is larger when individuals listen to sounds containing a regular pattern compared to sounds without a regular pattern (Pantev et al., 1994; Pantev et al., 1996; Gutschalk et al., 2002; Ross et al., 2002; Barascud et al., 2016; Sohoglu and Chait, 2016; Teki et al., 2016; Southwell et al., 2017; Herrmann and Johnsrude, 2018b; Southwell and Chait, 2018; Herrmann et al., 2019). Here, we investigated whether sustained neural activity changes when listeners learn a recurring regular pattern.

We observed that sustained activity increased earlier for recurring compared to novel patterns when participants attended to but not when they ignored the sounds (Figure 4). The earlier increase in sustained activity for recurring compared to novel patterns was present for trials within the first half of a stimulation block (Figure 5) and thus mirrors the pattern of response times observed in Experiment I (Figure 1; cf. Agus et al., 2010; Bianco et al., 2020). The earlier increase in sustained activity may thus index processes related to rapid perceptual learning.

### Reduced sustained activity due to pattern recurrence

We observed reduced sustained activity 1–2 s after pattern onset for recurring compared to novel patterns, both when participants actively or passively listened to sounds (Figure 4). This effect was present in the average response time courses of each of the four EEG experiments, but statistically significant only when data were pooled across experiments in our explorative analyses. This suggest that the reduced sustained activity for recurring compared to novel patterns is a reliable, but small effect, requiring high statistical power.

Our observation that sustained activity in the 1–2 s time window is reduced for recurring compared novel patterns regardless of participants’ attentional focus is in line with work showing that perceptual learning in adulthood, including learning of auditory patterns similar to the ones used here, can occur in the absence of attention or when patterns are task irrelevant (Seitz and Dinse, 2007; Andrillon et al., 2017; Bianco et al., 2020). Because recurring patterns are more predictable than novel patterns, the reduction in sustained activity for recurring relative to novel patterns is consistent with a predictive coding framework, in which responses to predictable events are reduced compared to responses to novel events (Friston, 2005; Baldeweg, 2006; Bubic et al., 2010; Arnal and Giraud, 2012; see also Heilbron and Chait, 2018). It remains an open question, however, whether reduced sustained activity reflects an attenuation of sensory signals as posited by a predictive coding framework (Friston, 2005; Friston and Kiebel, 2009) and consistent with suggested auditory cortex sources underlying sustained activity (Barascud et al., 2016; Herrmann et al., 2021), or whether reduced sustained activity for recurring patterns results from non-sensory prediction processes. Behavioral benefits for recurring over novel patterns could reflect successful prediction.

The reduction in sustained activity for recurring compared to novel patterns contrasts with results from other work using patterns in white noise stimuli and investigating inter-trial phase coherence in the 0.5 to 8 Hz frequency band (Luo et al., 2013; Andrillon et al., 2015). These studies report greater phase coherence for recurring patterns compared to novel patterns (Luo et al., 2013; Andrillon et al., 2015). It thus appears that reduced sustained activity and greater phase coherence may reflect perceptual learning independently.

The recurrence-related reduction in sustained activity 1–2 s post pattern onset was limited to late sound trials only (Figure 5), contrary to recurrence-related behavioral benefits that manifest after a few trials (Experiment I; Figure 1; cf. Agus et al., 2010; Bianco et al., 2020). These discrepant results suggest that reduced sustained activity for recurring compared to novel patterns may not be associated with the rapid perceptual learning observed behaviorally. We suggest that the recurrence-related reduction in sustained activity may instead be due to a learning mechanism that does not require attention and that enables subsequent processes, such as increased predictability or reduced novelty.

### Speculations about underlying neural sources

The underlying sources of pattern-related sustained activity involve auditory cortices (Pantev et al., 1994; Pantev et al., 1996; Gutschalk et al., 2002; Barascud et al., 2016; Herrmann et al., 2021) and potentially higher-level brain regions including the parietal cortex, frontal cortex, and hippocampus (Tiitinen et al., 2012; Barascud et al., 2016; Teki et al., 2016). Previous work suggests that there are two distinct regions, an anterior and a posterior one, in the auditory cortex that generate sustained activity, but that only the anterior region is modulated by auditory patterns (Gutschalk et al., 2002). Our 16-channel EEG setup does not allow firm conclusions to be made about the underlying sources of the learning-related neural effects, but the frontal-central EEG distribution depicted in Figure 4C is consistent with a more anterior region of auditory cortex as well as with frontal cortex activity. From the topographical EEG distribution, however, it appears unlikely that the parietal cortex is involved. Previous work suggests that perceptual learning recruits the hippocampus (Rose et al., 2011; Mundy et al., 2013; Larcombe et al., 2018), but whether or not the hippocampus contributes to long-term representations of auditory patterns and the sustained activity signatures thereof cannot be answered with our EEG setup. Other imaging modalities such as functional magnetic resonance imaging or lesion studies may be more suitable to assess the contribution of the hippocampus.

## Conclusions

In a series of five behavioral and EEG experiments, we investigated the behavioral and neural signatures that index rapid perceptual learning of regular patterns in sounds. We show that participants detect regular patterns faster and that they are more sensitive to pattern deviations and to the temporal order of pattern onset relative to a visual stimulus for recurring compared to novel regular patterns. Sustained neural activity indexed perceptual learning in two ways. Sustained activity increased earlier for recurring compared to novel patterns when participants attended to sounds, but not when they ignored them. This effect mirrored the rapid perceptual learning we observed behaviorally. The magnitude of sustained activity was reduced for recurring compared to novel patterns both when participants attended to, and ignored, the sounds. This reduction of sustained activity appeared only in later phases of our experimental protocols, suggesting it is not directly related to rapid perceptual learning, but perhaps due to a learning mechanism that does not require attention. Our study thus reveals neural markers of perceptual learning of auditory patterns, and of processes that may be related to reduced novelty or better prediction of learned auditory patterns.

## Acknowledgements

Research was supported by the Canadian Institutes of Health Research and Natural Sciences and Engineering Research Council grants to ISJ. BH was supported by a BrainsCAN postdoctoral fellowship (Canada First Research Excellence Fund; CFREF) and the Canada Research Chair program. We thank Ala Almanaseer and Mebaa Kassa for their help with data recording.

